# Stream–lake network position structures freshwater microbial communities across multiple catchments

**DOI:** 10.64898/2026.10.29.762755

**Authors:** Anish Kirtane, Kristy Deiner, Alex R. Hall

**Affiliations:** Department of Environmental Systems Science, Institute of Integrative Biology, ETH Zurich, Zurich 8092, Switzerland; Department of Environmental Systems Science, Institute of Biogeochemistry and Pollutant Dynamics, ETH Zurich, Zurich 8092, Switzerland; SimplexDNA AG, Winterthur, Switzerland 8404

**Keywords:** Freshwater microbiomes, bacterial biogeography, stream–lake continuum, hydrological connectivity, core microbiome, community turnover, 16S rRNA metabarcoding

## Abstract

Streams and lakes form hydrologically connected freshwater networks. Different environmental conditions and dispersal dynamics along the stream–lake continuum potentially affect bacterial community composition, although the strength of such effects may vary among individual catchments. Here, we analysed 224 water samples from 58 streams draining into eight lakes across Switzerland, covering the full continuum from headwaters to lake outlets. Bacterial community composition analyzed via 16S metabarcoding differed between streams and lakes, with a sharp turnover at stream-lake confluence, and this pattern was repeatable across all catchments. Lake communities had lower alpha diversity and lower overall beta dispersion than stream communities, despite hydrological connectivity with a diverse stream microbiome. Community composition in both lakes and streams differed between catchments, but this pattern was more pronounced in lakes. Lake communities were dominated by a small set of widespread and abundant core taxa, primarily established pelagic freshwater genera, that captured the catchment-level differences through shifts in relative abundance. In contrast, stream communities contained a comparatively less abundant core microbiome dominated by soil- and groundwater-associated genera, together with a larger and more diverse peripheral microbiome. Together, these findings show that bacterial community composition changes repeatably across the different positions of hydrological networks.

## Introduction

Over the past decades, numerous studies have shown that bacterial communities exhibit clear biogeographic patterns, including distance–decay relationships and associations with environmental gradients, indicating the joint influence of environmental selection and dispersal processes (Hanson et al., 2012; Martiny et al., 2006; Zeglin, 2015). The central debate in the field has shifted from whether bacteria exhibit biogeography to which forces govern community structure. Community assembly and metacommunity theory provide a conceptual framework for understanding the ecological processes that structure biological communities, including environmental selection (species sorting), dispersal limitation, mass effects, and stochastic colonization (Leibold et al., 2004; Vellend, 2010). The relative weight of deterministic and stochastic processes, and how they vary across spatial scales and ecosystem types, remains an active area of inquiry. Freshwater catchments can provide promising natural model systems for studying microbial community assembly, because environmental heterogeneity, dendritic network structure, and directional dispersal through hydrological flow create spatial and environmental gradients that may structure microbial communities.

Freshwater bacterial biogeography is of particular interest given that some important properties, such as nutrient removal, pathogen abundances, toxin production and heavy metal metabolism, are potentially influenced by bacteria in drinking water sources (Cotner & Biddanda, 2002; Hanson et al., 2012). Freshwater networks offer natural, discrete, replicated units for testing bacterial biogeography theory, each with multiple hydrological network positions (e.g., streams, lake inlets, pelagic lake waters, and lake outflows). Previous studies showed microbial communities change across connected stream–lake networks, but many have focused on individual networks, large regional gradients, or a limited number of systems (Crump et al., 2007; Niño-García et al., 2016; Papadopoulou & Lindström, 2026; Ruiz-González et al., 2015; Savio et al., 2015; Tang et al., 2020). Consequently, it remains unclear whether hydrological network position structures microbial communities consistently across independent catchments. Hydrological position can impact the physical and chemical environment a community experiences, primarily through increased water residence time in lotic positions (Niño-García et al., 2016; Read et al., 2015; Ruiz-González et al., 2015; Savio et al., 2015). Furthermore, hydrological network position can impact the dispersal corridors available for microbial taxa, for example via unidirectional flow from headwaters to lakes, making it a compound axis of both environmental filtering and connectivity (Besemer et al., 2013). Thus, hydrological network position may structure microbial communities in a consistent way, independent of the geographic location of the catchment, but testing this requires sampling across hydrological network positions in multiple replicated catchments, a framework rarely applied to freshwater microbial biogeography.

Sampling multiple catchments as replicates raises a second unresolved question: how does community structure vary across catchments within each hydrological context? Catchment identity represents the combined influence of multiple context-dependent factors, including geology, land use, climate, hydrology, nutrient regime, etc. (Diniz-Filho et al., 2012). These characteristics can alter environmental conditions and impose distinctive pressures on local microbial communities (Adams et al., 2010; Zeglin, 2015). This may lead to sites with more similar environmental conditions having more similar microbial communities. Furthermore, freshwater bacterial communities commonly exhibit distance–decay relationships, with more geographically distant sites having relatively dissimilar communities, potentially reflecting the influence of dispersal limitation or other drivers of community assembly independent of measured environmental conditions (Hanson et al., 2012; Martiny et al., 2006). Finally, differences in community composition among hydrological positions within catchments may also be modulated by individual catchment context. Thus, comparing community composition at multiple hydrological positions and in multiple catchments may reveal patterns that are consistent across catchments, or that are specific to individual catchments, and how they relate to spatial distance and environmental differences.

Freshwater bacterial communities often contain a very large number of taxa, but not all taxa contribute equally to observed biogeographic patterns. A common approach is to focus on taxa that are both prevalent and abundant within a habitat (i.e., the core microbiome) because they likely contribute key functions to the community (Neu et al., 2021; Risely, 2020). The core microbiome is often interpreted as reflecting environmentally filtered assembly, where taxa can persist reliably under a given set of environmental conditions. (Besemer et al., 2013; Neu et al., 2021; Tovar-Herrera et al., 2026). In contrast, the peripheral microbiome comprises low-prevalence taxa that are inconsistently detected across samples and contribute disproportionately to compositional turnover (Besemer et al., 2013; Gurung et al., 2025). Because core taxa are expected to reflect persistent responses to local environmental conditions, whereas peripheral taxa may be more strongly influenced by stochastic recruitment and dispersal, comparing their spatial patterns can provide insight into the relative importance of deterministic and stochastic assembly processes across environments.

Here, we employ a high-resolution water sampling design in eight catchments in Switzerland, collecting samples every few kilometers from the headwaters down to the lake outlets. We analyze the water samples primarily by 16S rRNA metabarcoding to address the following questions: 1) Does hydrological network position (streams vs lakes) structure microbial communities in consistent ways across catchments, independently of catchment identity? 2) Does the bacterial community composition vary among different catchments, and does the strength of this effect differ for streams and lakes? 3) Which members of the microbial community contribute most strongly to spatial differentiation across streams and lakes, and does this differ between catchments?

## Methods

### Sampling design, collection, and processing

We describe detailed field sampling methods in Kirtane et al. (2026). Briefly, we collected 224 water samples from 58 streams distributed across eight lake catchments in Switzerland (Figure 1B). Six of the eight lakes formed three pairs of hydrologically connected systems: Lake Pfaeffikon-Greifen, Hallwil-Baldegg, and Sils-Silvaplana.Within each catchment, we sampled sites from the headwaters downstream to the lake and its outlet and assigned each site to one of four hydrological network position categories (Figure 1A): Up-stream, Inlet, Lake, and Outflow. We classified all stream samples as “Up-stream,” except for the final sampling point immediately upstream of the stream–lake confluence, which we classified as “Inlet.” We collected “Lake” samples from the lake surface within 5–50 m of the stream–lake confluence. We collected “Outflow” samples from the stream channel downstream of the lake outlet, within 100 m of where water exited the lake. Detailed maps of sample locations are provided in Kirtane et al. (2026). Within each catchment, we completed all sampling within a few days to minimize temporal variability (Table 1).

**Figure 1:**
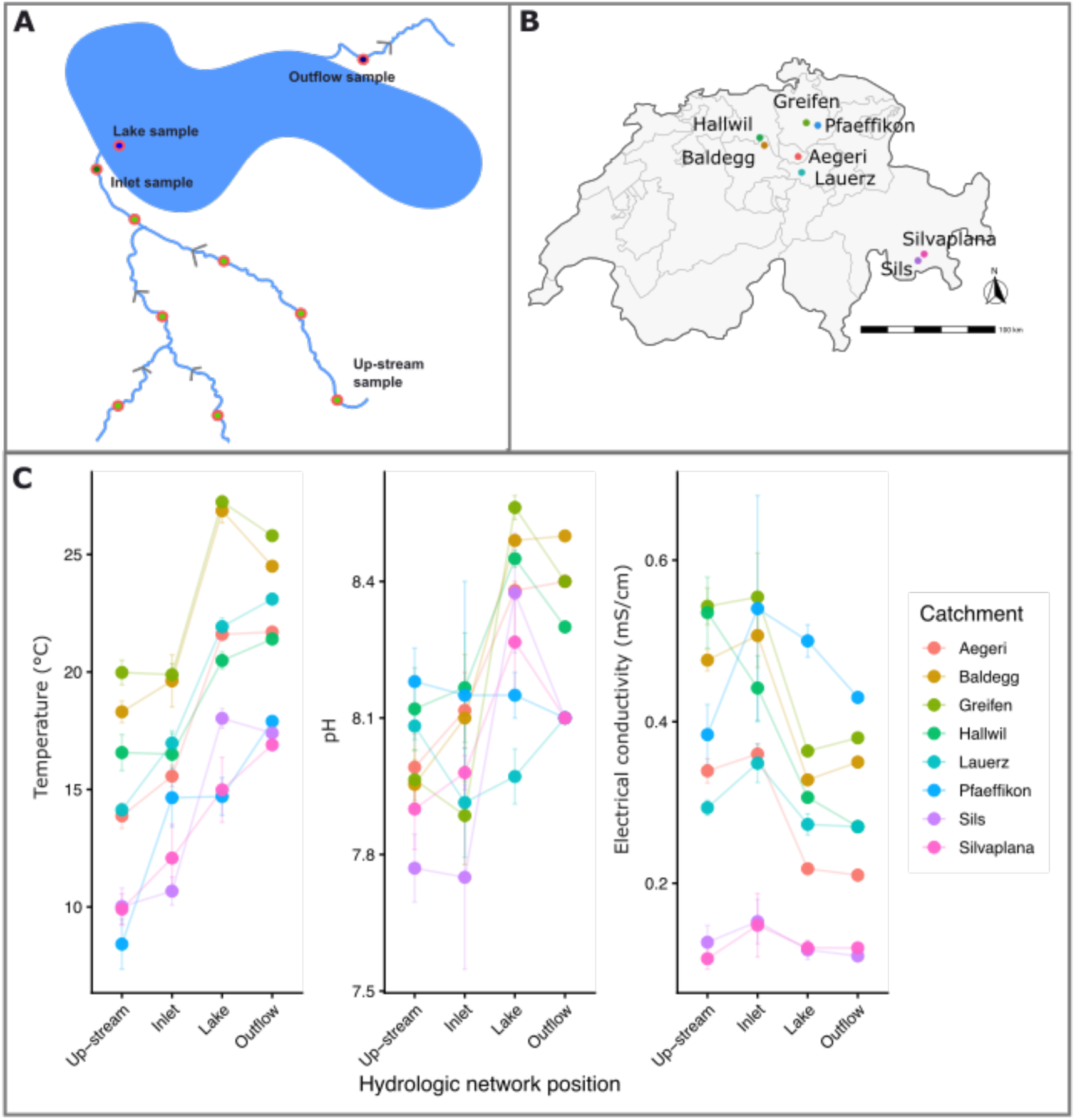
(A) Simplified illustration of sampling design within each catchment. Red dots indicate sampling locations, and grey arrows indicate the direction of water flow (adapted from Kirtane et al., (2026), licensed under CC BY-NC 4.0). (B) Geographic distribution of sampled catchments across Switzerland (adapted from Kirtane et al., (2026), licensed under CC BY-NC 4.0). Maps of sampling locations in each catchment can be found in Kirtane et al., (2026). (C) Physicochemical trajectories along the hydrological gradient for each catchment. Points show mean temperature (°C), pH, and electrical conductivity (mS/cm) at each network position, with lines connecting sites along the flow path. Error bars represent ± standard error (SE) of replicate samples. Data for this plot are from Kirtane et al., 2026.

**Table 1:**
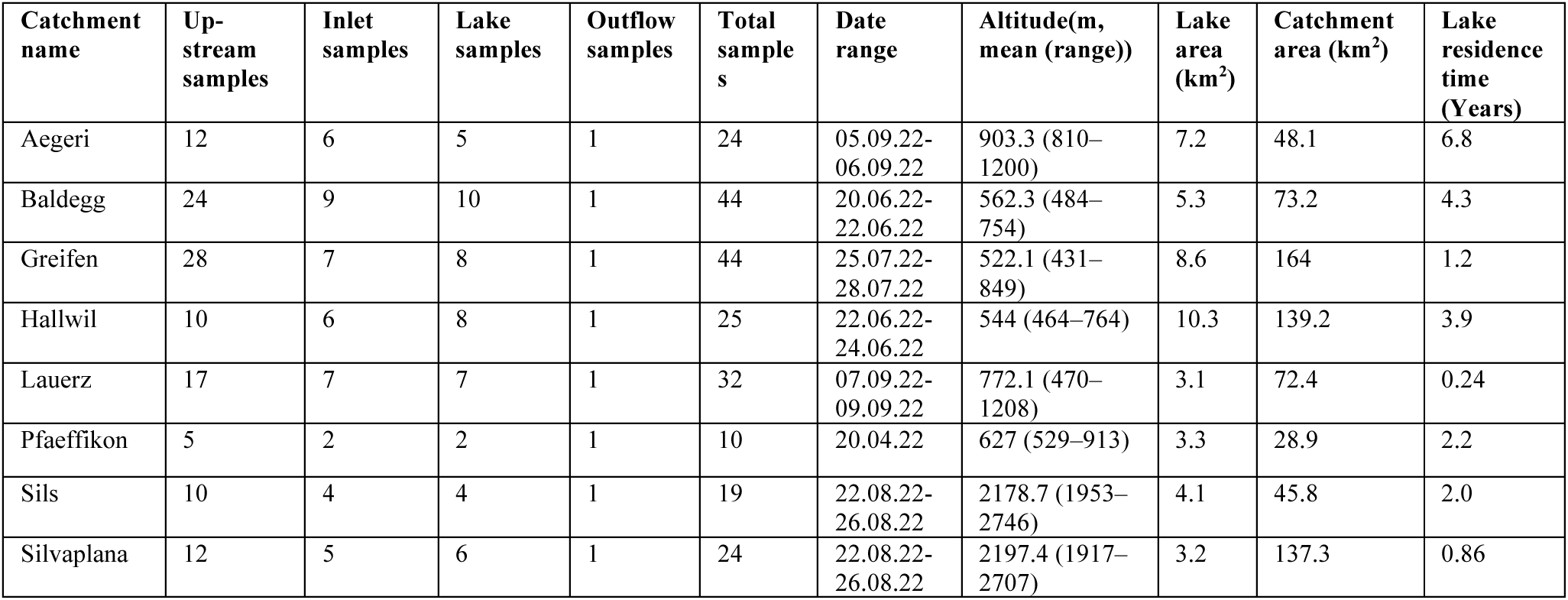
Overview of sampling effort and key characteristics of the eight lake catchments, including the number of samples collected at each hydrological position, altitude range, lake area, and catchment area.

| Catchment name | Up-stream samples | Inlet samples | Lake samples | Outflow samples | Total samples | Date range | Altitude(m, mean (range)) | Lake area (km<sup>2</sup>) | Catchment area (km<sup>2</sup>) | Lake residence time (Years) |
| --- | --- | --- | --- | --- | --- | --- | --- | --- | --- | --- |
| Aegeri | 12 | 6 | 5 | 1 | 24 | 05.09.22-06.09.22 | 903.3 (810–1200) | 7.2 | 48.1 | 6.8 |
| Baldegg | 24 | 9 | 10 | 1 | 44 | 20.06.22-22.06.22 | 562.3 (484–754) | 5.3 | 73.2 | 4.3 |
| Greifen | 28 | 7 | 8 | 1 | 44 | 25.07.22-28.07.22 | 522.1 (431–849) | 8.6 | 164 | 1.2 |
| Hallwil | 10 | 6 | 8 | 1 | 25 | 22.06.22-24.06.22 | 544 (464–764) | 10.3 | 139.2 | 3.9 |
| Lauerz | 17 | 7 | 7 | 1 | 32 | 07.09.22-09.09.22 | 772.1 (470–1208) | 3.1 | 72.4 | 0.24 |
| Pfäeffikon | 5 | 2 | 2 | 1 | 10 | 20.04.22 | 627 (529–913) | 3.3 | 28.9 | 2.2 |
| Sils | 10 | 4 | 4 | 1 | 19 | 22.08.22-26.08.22 | 2178.7 (1953–2746) | 4.1 | 45.8 | 2.0 |
| Silvaplana | 12 | 5 | 6 | 1 | 24 | 22.08.22-26.08.22 | 2197.4 (1917–2707) | 3.2 | 137.3 | 0.86 |

At each site, we collected 100 mL of water and filtered it through a 0.22 μm Isopore polycarbonate filter (GTTP02500; Millipore) mounted in a 25 mm Swinnex filter holder using 50 mL syringes. We processed samples following the workflow described previously (Kirtane et al., 2026). In this study, however, we only analyzed the membrane-bound state because previous studies showed that this state often contains the highest diversity and is most commonly examined in microbial ecology (Kirtane et al., 2023, 2026; Nagler et al., 2022). To provide a physicochemical context, we used metadata (temperature, pH, and electrical conductivity) collected at each sampling site in Kirtane et al. (2026).

### 16S rRNA Metabarcoding

We conducted 16S rRNA metabarcoding on a total of 224 previously extracted DNA samples. We also included 22 negative controls at the DNA extraction and PCR steps to identify contamination. We used a two-step PCR approach, where PCR1 involved amplification of the 16S rRNA target region with 25 cycles, followed by magnetic bead clean-up, and PCR2 involved 10 cycles that amplified the initial PCR1 amplicon while adding dual-unique indexes and Illumina adapters.

We used the following 16S primers (341F: 5’-CCTACGGGNGGCWGCAG-3’; 806R: 5’- GGACTACNVGGGTWTCTAAT-3’) spanning the V3–V4 region, becuase in-silico analysis showed this set to capture the largest amount of bacterial diversity from river and lake samples (Zhang et al., 2023). We pooled all four variants of the base-shifted primers at 50 µM concentration. PCR1 reactions were run in triplicate in 10 µL volumes containing 5 µL KAPA HiFi HotStart ReadyMix (Roche), 0.1 µL of 50 µM primer mix, 3.4 µL of water, and 1.5 µL of template DNA. PCR1 consisted of an initial incubation at 95 °C for 3 min, followed by 25 cycles of denaturation at 95 °C for 30 s, annealing at 50 °C for 30 s, and extension at 72 °C for 30 s, with a final extension at 72 °C for 5 min. Triplicates were pooled and cleaned using Ampure beads at a 0.9:1 bead-to-PCR product ratio.

We performed PCR2 using Illumina UDI 10 nt index primers (Set A, Plates 1–5; Illumina) to enable unique dual indexing for multiplexed sequencing. Each reaction contained 12.5 µL KAPA HiFi HotStart ReadyMix, 2.5 µL of 10 µM index primers, 3 µL of cleaned PCR1 product, and 7 µL of water (25 µL total). PCR2 consisted of an initial incubation at 95 °C for 3 min, followed by 10 cycles of denaturation at 95 °C for 30 s, annealing at 55 °C for 30 s, and extension at 72 °C for 30 s, with a final extension at 72 °C for 5 min. PCR2 products were cleaned using Ampure beads at a 0.9:1 ratio.

We quantified the cleaned PCR2 product using a Spark® 10M Multimode Microplate Reader (Tecan, Männerdorf, Switzerland) with Qubit BR Kit reagents, analyzed in triplicate, and normalized using a pipetting robot (Liquid Handling Station, BRAND Scientific, Essex, CT, USA). The pooled library was subjected to a reconditioning PCR (12.5 µL KAPA HiFi ReadyMix, 1 µL each of P5 and P7 primers, 3 µL pooled library, and 6.5 µL water), consisting of an initial incubation at 95 °C for 3 min, followed by 4 cycles of denaturation at 95 °C for 30 s, annealing at 60 °C for 30 s, and extension at 72 °C for 30 s, with a final extension at 72 °C for 5 min. The resulting library was cleaned with Ampure beads and analyzed using a D1000 HS ScreenTape Assay Kit on a 4150 TapeStation (Agilent Technologies, Santa Clara, CA) to confirm removal of small DNA fragments.

We sequenced the amplicon libraries on the AVITI System (Element Biosciences) using the Cloudbreak™ 2×300 bp Sequencing Kit (Medium Output), generating paired-end reads with a yield of approximately 300 million reads at the Lausanne Genomic Technologies Facility (Lausanne, Switzerland), with 20% PhiX added prior to sequencing.

### Data analysis

We removed PhiX contamination and low-complexity reads, trimmed ends, and merged pairs using ‘USEARCH v11.0.667’s (min overlap = 30 bp, min %id = 60, max mismatches = 20, min merged length = 100, no minimum quality filter during merging). We filtered reads based on primer matching, retaining sequences with ≤2 mismatches that yielded amplicons of 200–600 bp. Reads with 3′-end mismatches or incorrect orientation were excluded, and full primer sequences were trimmed. We applied post-merging filters with ‘PRINSEQ-lite v0.20.4’ (length 150–550 bp, GC 30–70%, mean Q ≥ 20, no ambiguous bases, DUST = 30 for low-complexity removal). After the bioinformatics filtration, we further cleaned the dataset in the phyloseq environment by removing amplicon sequence variants (ASVs) that lacked a Phylum-level assignment, ASVs with fewer than 500 total reads, and ASVs that occurred in only one sample (McMurdie & Holmes, 2013). We also removed ASVs flagged as contaminants in the negative controls using prevalence-based contamination detection in the ‘decontam’ package (threshold = 0.1; Davis et al., 2018). Taxonomy was assigned to ASVs using the SILVA reference database (version 138.1; Quast et al., 2012). We conducted all data analyses in R 4.4.2 (R Core Team, 2024) using ‘phyloseq’ (McMurdie & Holmes, 2013) and ‘tidyverse’(Wickham et al., 2019) for most analyses. We set the significance threshold at α = 0.05 unless stated otherwise. All samples were rarefied to 100,000 reads per sample, which retained 222 of 224 samples while capturing 98.8% of the observed ASV richness. After data cleaning and rarefaction, the resulting dataset contained 34,938 ASVs, with taxonomic assignments at 100% for Kingdom and Phylum, 99.1% for Class, 91.5% for Order, 82.3% for Family, and 61.1% for Genus.

Physicochemical variation was analysed using linear mixed-effects models with the ‘lme4’ package (Bates et al., 2015), testing hydrological network position as a fixed factor and including catchment identity as a random intercept and random slope to account for non-independence among samples within the same catchment. A second analysis, including stream identity as a nested random effect to account for repeated sampling within streams, did not substantially alter qualitative conclusions (Table S3). Significance of fixed effects was tested using Satterthwaite’s approximation for degrees of freedom, implemented in the ‘lmerTest’ package (Kuznetsova et al., 2017). We tested for differences in alpha diversity among hydrological network positions using a similar linear model, with observed ASV richness as the response variable, hydrological network position as a fixed effect, and catchment as a random intercept and random slope. Richness was log-transformed to meet model assumptions. The same model structure was applied to Shannon diversity (without log transformation). For post hoc pairwise comparisons among hydrological positions, we used estimated marginal means with Tukey-adjusted p-values to account for multiple testing. Including stream as a nested random effect did not change the results (Table S3). To assess whether average alpha diversity and physicochemical conditions differed among catchments independent of hydrological position, we fitted additive linear models using catchment and hydrological position as fixed effects to mean values calculated for each hydrological position within each catchment.

To analyse differences in bacterial community composition among samples, we calculated Bray–Curtis dissimilarities at the ASV level and visualized them using principal coordinates analysis (PCoA). We tested the effect of hydrological network position using PERMANOVA with permutations constrained within catchments to account for the hierarchical sampling structure. We then evaluated the homogeneity of multivariate dispersion among hydrological network positions using PERMDISP with permutation-based ANOVA and Tukey-adjusted pairwise comparisons. To investigate differences among catchments, we divided the dataset into stream (up-stream and inlet) and lake (lake and outflow) communities and performed all subsequent analyses separately for each subset. We tested for differences among catchments using PERMANOVA with catchment specified as the explanatory variable. Finally, we repeated these analyses using both Bray–Curtis and Jaccard distance matrices to determine whether the observed patterns were consistent when considering taxon abundances or taxon occurrence alone.

We tested for an association between geographic distance (km), altitude (m), and sampling-date differences (days) with dissimilarity of community composition (Bray-Curtis dissimilarity) among pairs of samples using Mantel tests (Pearson correlation, 999 permutations) with the ‘vegan’ package (Dixon, 2003). To evaluate catchment-scale patterns, Mantel tests were performed using only between-catchment pairs of samples. We further used Mantel tests to test associations between community-composition dissimilarity and environmental dissimilarity, first testing three variables (temperature, electrical conductivity, and pH) individually, and then as an aggregate multivariate measure (Euclidean distance between each pair of samples across the three variables, after rank transformation).

To partition communities into core and peripheral microbiota, we agglomerated ASVs at the genus level. Core genera were defined using a common core concept (Risely, 2020) with combined occurrence-abundance thresholds (Neu et al., 2021). Genera with a relative abundance of at least 0.1% in more than 90% of samples and occurring in all eight catchments were classified as core; all remaining genera were classified as peripheral. Preliminary analysis evaluating how the number of core genera varies across prevalence (60– 99%) and relative abundance thresholds (0.01–1%) showed strong sensitivity to threshold choice (Figure S1). The selected thresholds (90% prevalence, 0.1% abundance) were chosen as a pragmatic compromise between permissive thresholds that inflate the core with low-abundance genera and stringent thresholds that reduce the core to very few taxa, limiting the ability to detect differences in community assembly across habitats (Figure S1). To determine which components of the community underlie the previously observed catchment-level structuring, we repeated the PERMANOVA analysis separately for the full, core, and peripheral communities using Bray–Curtis dissimilarities calculated from genus-level relative abundance matrices for each habitat. Relative read abundance was calculated for each sample by dividing genus-level read counts by the total read count per sample. Mean relative read abundance (%) was then computed per genus within each catchment and habitat. For visualization purposes, core genera contributing <3% mean relative read abundance within a catchment were grouped as ‘Other Core’ to emphasize dominant taxa.

To assess whether bacteria characteristic of lake communities were also detected upstream, we first selected genera that met the core criteria in lake samples but not in stream samples. For each of these lake-core-only genera, and for each catchment separately, we identified ASVs detected in lake samples and then assessed their occurrence in upstream samples from the same catchment. We excluded lake inlets here. We considered ASVs from lake-core-only genera as detected in a given upstream sample if there were at least five reads. For each lake-core-only genus and in each catchment, we then calculated 1) the percentage of ASVs also detected upstream in at least one sample and 2) the percentage of upstream samples from the same catchment that contained at least one ASV from a lake-core-only genus. We visualized the results using a bubble matrix plot.

## Results

### Hydrological network position structures bacterial community composition

Microbial diversity, taken as observed ASV richness or Shannon diversity, varied among water samples from different hydrological positions (Figure 2A; Table S1). Diversity was highest in upstream and inlet sites, which did not differ significantly from one another (Observed: p = 0.974; Shannon: p = 0.860) (Table S2). In contrast, lake and outflow sites had lower diversity than upstream sites on average (all p ≤ 0.0026 for observed richness; all p ≤ 0.0008 for Shannon diversity) (Table S2). Lake and outflow positions did not differ significantly for either diversity metric (Observed: p = 0.909; Shannon: p = 0.991; Table S2). This pattern was consistent across the eight catchments, although average alpha diversity differed among catchments for both observed richness (two-way ANOVA, F_7,21_ = 3.58, p = 0.011) and Shannon diversity (two-way ANOVA, F_7,21_ = 6.183, p < 0.0011; Figure 2A). Environmental variables showed parallel gradients across hydrological positions, with lake samples being warmer (linear mixed-effects model: F_3,9.87_ = 58.45, p < 0.001), more alkaline (F_3,12.11_ = 5.48, p = 0.013), and less electrically conductive (F_3,9.14_ = 7.16, p = 0.009) than stream samples (Figure 1C). As for diversity, there were average differences among catchments in terms of temperature and conductivity (ANOVA: temperature, p < 0.001; conductivity, p < 0.001), but not pH (p = 0.065; Figure 1C).

**Figure 2:**
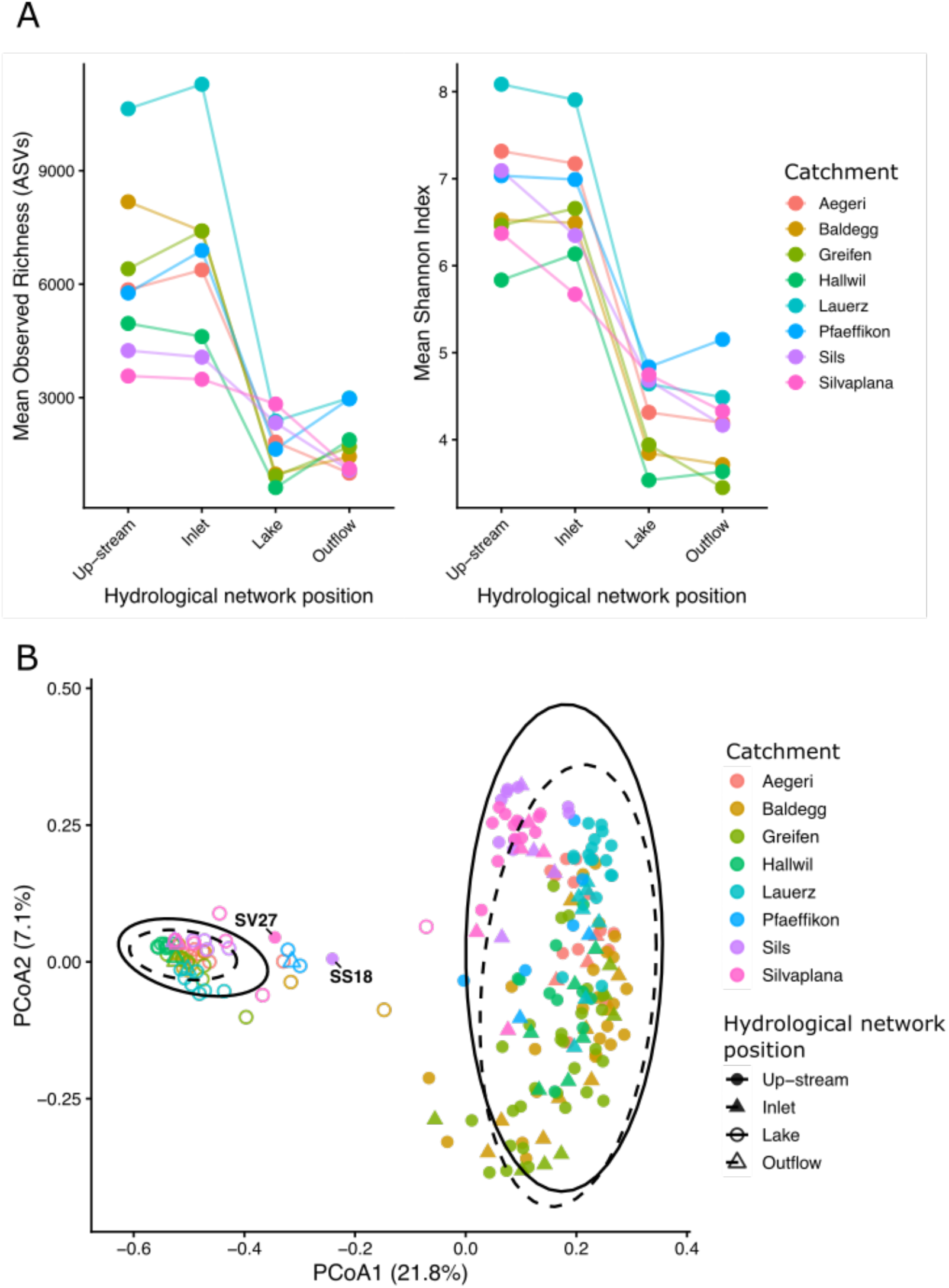
(A) Patterns of alpha diversity along the hydrological network gradient across eight catchments. Points represent mean observed ASV richness (left panel) and mean Shannon diversity (right panel) at each hydrological position (Up-stream, Inlet, Lake, Outflow) within each catchment, with lines connecting sites along the flow path; data for individual samples are shown in Figure S4. (B) PCoA plot of Bray–Curtis dissimilarities illustrating patterns of community composition across hydrological network positions and catchments. Points represent individual samples, colored by catchment and shaped according to hydrological position (see legend). Ellipses represent the 95% dispersion of samples around group centroids. Two up-stream samples, SS18 from Sils and SV27 from Silvaplana, that cluster together with the lake and outflow samples, are labelled.

Differences among hydrological positions were also evident when we examined beta diversity. Up-stream and inlet samples clustered separately from the lake and outflow samples on the PCoA plot (Figure 2B, based on Bray-Curtis distances). This clustering was supported by a PERMANOVA with permutations restricted within catchments, indicating that hydrological network position explained 21.2% of the variation in community composition (Bray–Curtis PERMANOVA, F_3,218_ = 19.60, R² = 0.212, p = 0.001, Table S4). Two stream samples clustered with lake communities (Figure 2B). These were from Sils and Silvaplana, and upon closer inspection of their geographic location, we concluded these streams were the outflows from small glacial lakes (Figure S3). Therefore, we excluded them from subsequent analyses to ensure consistency between sample classification and the hydrological framework underlying the study design. The PCoA plot shows lake and outflow samples cluster relatively tightly together, whereas upstream and inlet samples are more dispersed in ordination space. This pattern was confirmed by PERMDISP (F₃,₂₁₈ = 78.19, p < 0.001; Figure S2, Tables S5–S6). This dispersion reflects differences in community composition among samples across all catchments, including within- and among-catchment differences. Together, these results indicate that lake and outflow samples were different and less variable in terms of community composition compared with upstream and inlet samples.

### Lakes show stronger between-catchment differentiation than streams

We tested for differences among catchments separately for stream (upstream and inlet) and lake (lake and outflow) samples, because of the clear separation of these groups of samples observed above. The lake communities clustered by catchment (Figure 3A). This was corroborated by PERMANOVA showing over 72% of the variance in microbial community (ASVs) composition among lake samples was explained by catchment identity (PERMANOVA, Bray–Curtis, R² = 0.72, F_7_ = 18.80, p = 0.001). Using an alternative dissimilarity measure revealed a similar qualitative trend, albeit with a lower proportion of variance explained (PERMANOVA using Jaccard distance instead of Bray-Curtis, R² = 0.39, F_7_ = 4.56, p = 0.001). Because Jaccard distance is sensitive to which ASVs are detected, but not their relative abundances, whereas Bray-Curtis distance is sensitive to both, these analyses indicate both types of dissimilarity contributed to differences among catchments in terms of lake community composition. Hydrologically connected lakes (e.g., Hallwil and Baldegg; Sils and Silvaplana) clustered more closely than other lakes. An exception to this trend was Lake Pfaeffikon, which clustered closer to high-elevation lakes, Sils and Silvaplana, despite being hydrologically connected to Lake Greifen (Figure 3A, Table 1).

**Figure 3:**
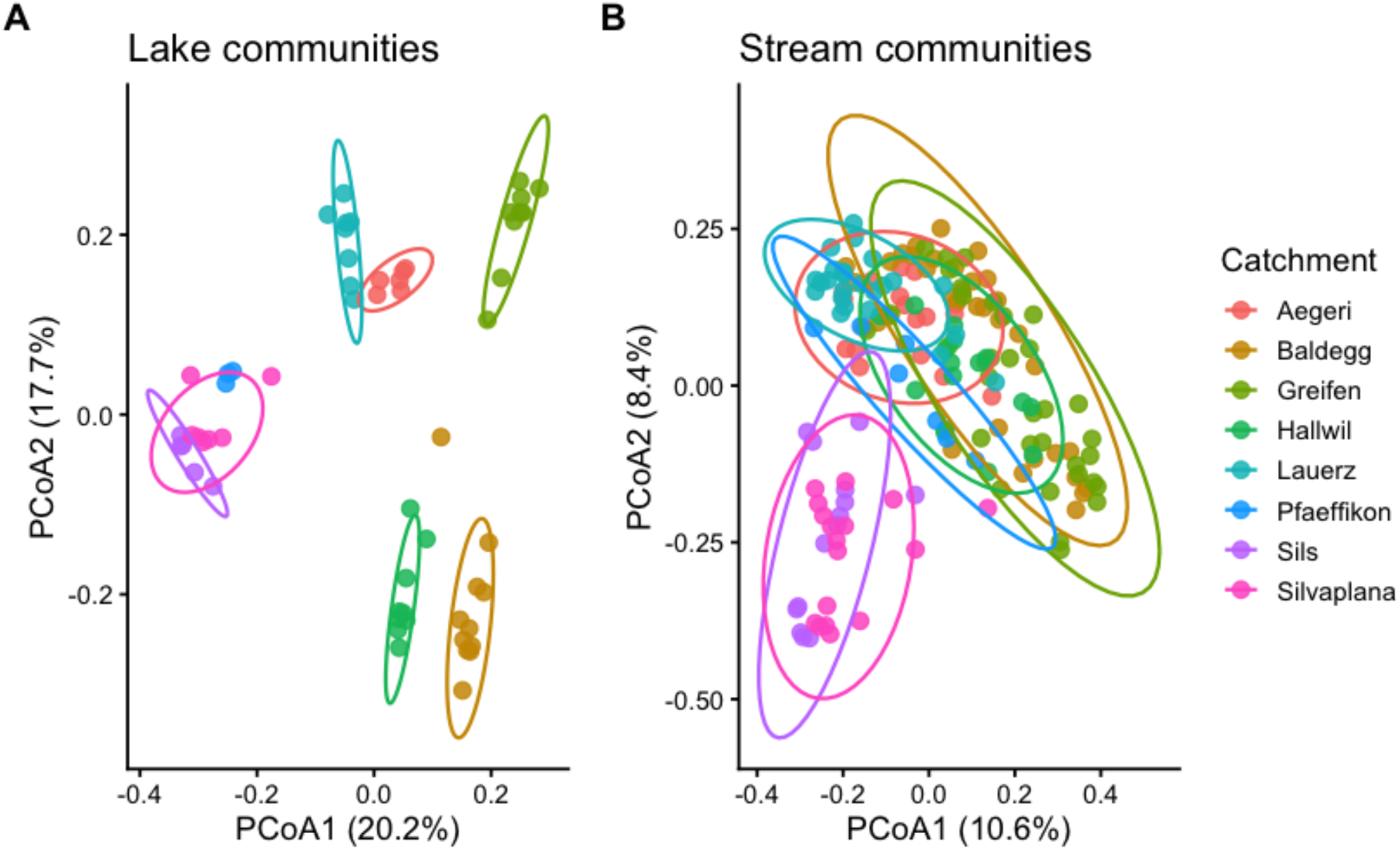
PCoA ordination of Bray–Curtis distances showing distinct patterns of community composition among catchments in lake (A) and stream (B) habitats. Samples are colored by catchment, and ellipses represent the 95% dispersion of samples around group centroids.

Although stream communities were, in terms of community composition across the entire dataset, more dispersed overall (Figure 2B, S2), stream samples from different catchments formed visibly separate clusters to a lesser extent than lake samples (Figure 3B). Nevertheless, catchment identity did explain a significant proportion of the variance of stream community composition (ASVs) (PERMANOVA, Bray–Curtis, R² = 0.21, F_7_ = 5.93, p = 0.001). A similar pattern was observed using Jaccard distances (PERMANOVA, Jaccard, R² = 0.20, F_7_ = 5.57, p = 0.001). This suggests differences in terms of which ASVs were detected, even without accounting for their relative abundances, explained much of the observed variation among stream communities from different catchments. The most obvious catchment-level clustering among stream samples was for those from Sils and Silvaplana, which clustered together and separately from other catchments, both in this ordination taking only stream samples (Figure 3B), and above with all hydrological positions together (Figure 2B). The differences among catchments remained significant when Sils and Silvaplana were excluded from the analysis (PERMANOVA, Bray–Curtis, R² = 0.17, F_5_ = 5.20, p = 0.001), indicating additional catchment-level differences between other locations. Unlike lake samples, Pfaeffikon stream samples did not cluster with samples from high-elevation streams in Sils and Silvaplana catchments (Figure 3B).

### Community structure varies with geographic and environmental differences

Community dissimilarity between pairs of samples increased significantly with both geographic distance and altitude differences in both streams and lakes (Table S7). This indicates samples from geographically and topographically more similar sites harboured more similar microbial communities. The Mantel tests included only between-catchment sample pairs. Catchment identity was strongly associated with community composition (Figure 3), and samples within the same catchment were generally both geographically close and compositionally similar. Including these within-catchment pairs could therefore have inflated the association between geographic distance and community dissimilarity. Like community dissimilarity, aggregate environmental dissimilarity (based on temperature, EC, and pH) was significantly associated with both geographic distance and altitude (Table S8), with temperature and EC driving most of this pattern (Table S7). Finally, environmental dissimilarity was significantly correlated with community dissimilarity (Table S8). Thus, sampling locations in different catchments that were further apart in space were also more dissimilar in both their recorded environmental conditions and microbial community composition, and this pattern was observed for both lake and stream samples.

Catchments were sampled on different dates, whereas all samples within each catchment were collected within a few days (Table 1). Thus, temporal effects could not be separated fully from catchment effects. Community dissimilarity between sample pairs from different catchments was positively associated with sampling-date differences in lakes (Mantel test, *p* = 0.01, *r* = 0.246), though not in streams (Mantel test, *p* = 0.971) (Table S8). Thus, we cannot exclude that sampling date contributed to some among-catchment differences, for example, via seasonal changes in environmental conditions (Eiler et al., 2012; Stadler & Del Giorgio, 2022). In line with this, aggregate environmental dissimilarity was positively correlated with sampling-date differences in both streams (Mantel *r* = 0.151) and lakes (*r* = 0.114) (Table S8, Figure S5). Despite this, the following observations suggest sampling date effects are unlikely to explain our main results. First, comparisons among hydrological network positions within catchments are unlikely to be impacted by sampling date, because all sites within each catchment were sampled within only a few days (Table 1). Second, differences between lake and stream communities were consistent across catchments, even though catchments were sampled on different dates (Figure 2). Third, there was no obvious trend for alpha diversity to change directionally with sampling date (Figure S5), suggesting our ability to detect diverse communities was not specific to early/late samples. Finally, because geographically proximate locations were generally sampled during the same excursions for logistical reasons, the association between community dissimilarity and sampling-date differences could also reflect shared geographic and environmental conditions. This explanation cannot be separated from seasonal change in our dataset.

### Lake microbiomes are dominated by a shared core assemblage

Core microbiome genera were defined separately for stream and lake habitats as genera exceeding 0.1% relative read abundance within individual samples, present in more than 90% of samples within that habitat, and detected in all catchments; all remaining genera were classified as peripheral genera. This identified 14 core genera from the lake habitats and 26 core genera in the stream habitats, with only 4 shared between the two (Figure 4A). The core microbiome dominated lake communities, accounting for 80.6% of total read abundance, whereas they contributed only 41.1% in streams (Figure 4B). Catchment-level structuring, which was stronger in lakes than in streams (Figure 3), was consistent across both core and peripheral microbiomes (Table S9).

**Figure 4:**
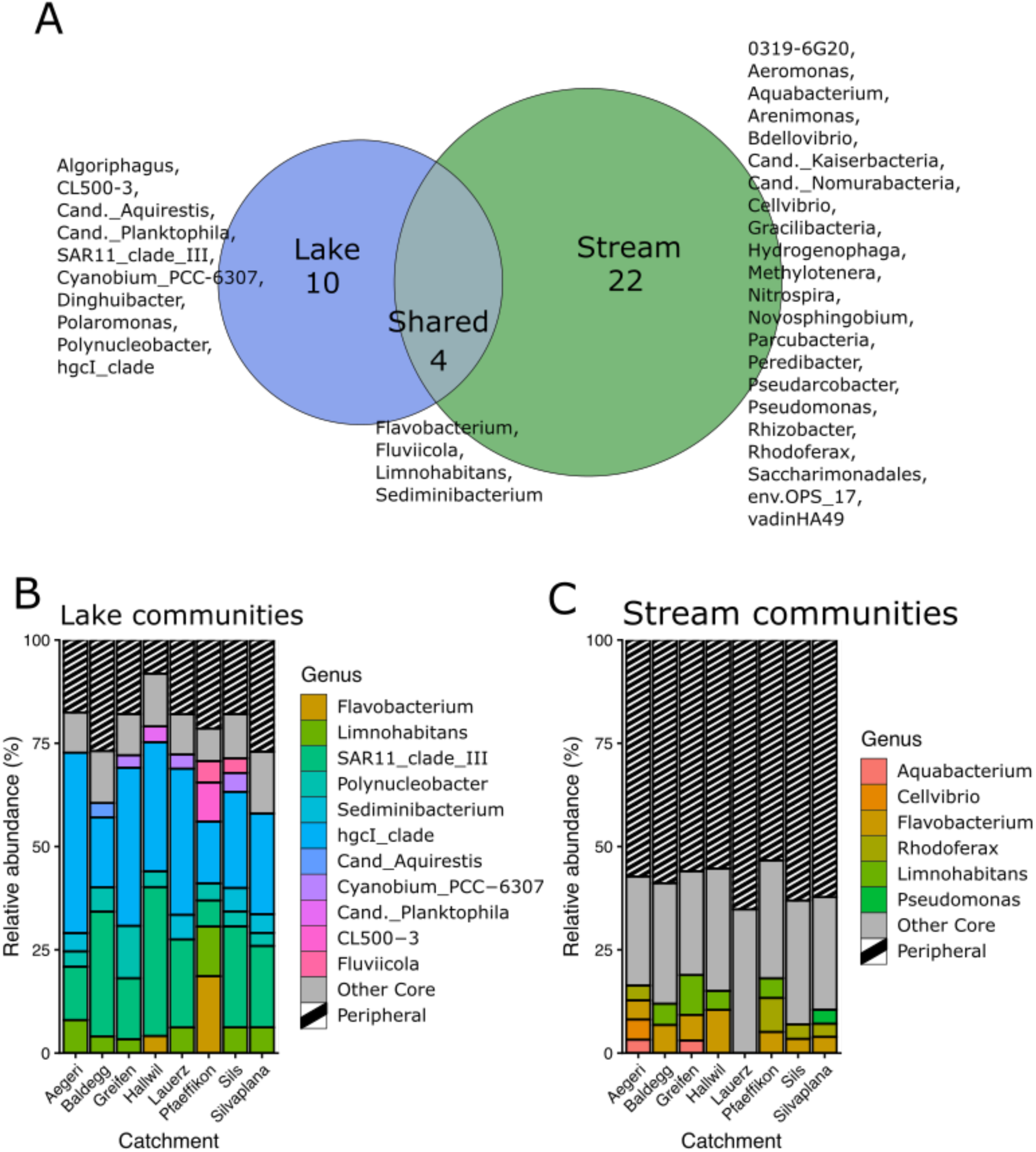
(A) Venn diagram showing unique and shared core genera between lake and stream communities. (B, C) Stacked bar plots showing mean relative read abundance (%) of dominant core genera across catchments in (B) lake and (C) stream communities. Core genera contributing <3% were grouped as ‘Other Core’.

Lake core microbiomes were dominated by well-established freshwater bacterioplankton lineages, including SAR11 clade III (LD12), *Polynucleobacter*, *Limnohabitans*, and members of the Actinobacteria (eg. hgcI clade) (Figure 4B). These taxa have been repeatedly identified as cosmopolitan freshwater specialists through cultivation, genomic, and global 16S surveys (Hahn, 2003; Hahn et al., 2012; Newton et al., 2011; Savio et al., 2015; J. Wang & Jia, 2016; Zaremba-Niedzwiedzka et al., 2013). Their consistent dominance across catchments supports the interpretation that lake biogeography is deterministically structured around a shared pelagic freshwater assemblage (Eiler et al., 2012; Lindström et al., 2005). All ten genus-level groups that met the core criteria in lakes but not in streams contained ASVs that were, in at least some catchments, shared between lake and upstream samples within the same catchment (Figure S6). Such overlap occurred in all eight catchments for *Polaromonas*, *Polynucleobacter*, and *hgcI clade*, and in four to seven catchments for the remaining genera. Lake inlet samples were excluded from this comparison. In contrast, stream cores included nitrifying and redox-associated taxa such as *Nitrospira* (Daims et al., 2015), freshwater methylotrophs such as *Methylotenera* (Kalyuzhnaya et al., 2012), soil-associated genera including *Cellvibrio* (DeBoy et al., 2008), and groundwater-linked candidate phyla radiation lineages such as *Parcubacteria* (Brown et al., 2015) (Figure 4B).

## Discussion

Our results quantify variation of bacterial community structure in relation to key potential contributing factors within and between catchments. Across all eight catchments, hydrological network position consistently structured bacterial communities, with the strongest compositional shift occurring at the stream-lake confluence. At the same time, community composition differed among catchments in both streams and lakes, although these differences were more pronounced in lakes. This stronger spatial structuring in lakes was primarily associated with shifts in the relative read abundance of core taxa, whereas stream communities were characterized by a larger contribution of peripheral taxa and higher compositional turnover among samples. Differences among catchments reflected broader spatial and environmental gradients. Catchments that were more similar in their geographic location, altitude, and measured environmental conditions also tended to harbour more similar bacterial communities. Together, these findings suggest catchment-scale variation in bacterial communities is structured by underlying environmental and spatial processes.

Our finding of pronounced bacterial community turnover across stream-lake confluences is consistent with observations made in previous studies (Crump et al., 2007; Kirtane et al., 2026; Niño-García et al., 2016; Papadopoulou & Lindström, 2026; Ruiz-González et al., 2015; Tang et al., 2020). By replicating across multiple confluences spanning diverse physicochemical conditions and sampling dates, we show that this pattern of bacterial turnover is repeatable within just 5–50 m of the stream–lake confluences. A comparable shift towards dominance by freshwater bacterioplankton was observed along the Danube River, where LD12, acI lineages, and *Polynucleobacter* increased downstream (Savio et al., 2015). Their prominence in our lake communities suggests that similar assemblages can develop both across stream–lake transitions and gradually along large rivers. In our study, all ten genera belonging to the lake core but not the stream core contained ASVs detected in both lake and upstream samples within the same catchment (Figure S6), indicating that members of these lake-associated groups were already present upstream. Their greater prominence in lakes is consistent with environmental filtering that favors a characteristic subset of the incoming microbiota. This interpretation is further supported by community convergence among lakes. For instance, a sample from Lake Hallwil was compositionally more similar to Lake Silvaplana, 140 km away, than to a directly inflowing stream only a few meters away.

What possible mechanisms explain the separation between lake and stream communities? One possible explanation involves physical transport processes, such as rapid dilution of stream communities or settling out of the water column upon flowing into the lakes (Kirtane et al., 2026; Niño-García et al., 2016; Papadopoulou & Lindström, 2026; Read et al., 2015). Such processes may be reinforced by thermal stratification at the stream-lake interface. The colder stream water (Figure 1C) may form a subsurface plume under the warmer lake water, physically disconnecting stream-transported bacteria from the lake surface (Kirtane et al., 2026). Support for the importance of transport and persistence processes also comes from parallel analyses of non-bacterial eDNA. Patterns of sharp turnover at the stream–lake confluence were also observed in the eDNA of plants and metazoans from the same samples (Kirtane et al., 2026). Given that eDNA from plants and metazoans is likely dominated by non-living material, this similarity may reflect the influence of transport and persistence processes rather than solely active community assembly. However, the current analysis cannot differentiate between living and dead bacteria, which may contribute to the high diversity of the peripheral microbiome through relic cells, despite isolating the membrane-bound state for analysis (Kirtane et al., 2026; Lennon et al., 2018). A second possible explanation is that different physicochemical conditions in lakes and streams select for different taxa. Consistent with this, the physicochemical conditions we measured differed between streams and lakes, with lake samples showing lower variability than stream samples (Figure 1C). The greater similarity of lake environments may therefore contribute to the convergence of bacterial communities across geographically distinct lakes through environmental filtering, whereby similar environmental conditions select for similar bacterial taxa. The presence of distinct core bacterial taxa in lakes further supports the idea that lake conditions consistently favour a characteristic subset of taxa. Together, these observations suggest that both physical transport processes and environmental filtering likely contribute to the pronounced turnover observed at stream-lake interfaces. Clarifying the balance between these processes is an important direction for future research (URycki et al., 2024).

In streams, the higher alpha diversity, larger relative proportion of peripheral taxa, and higher **overall** beta dispersion **(including within- and between-catchment differences)** of bacterial community composition may reflect multiple coacting assembly processes, including stochastic recruitment, particle-associated taxa, and specialization on narrower ecological niches compared to lake habitats (Hanson et al., 2012; Shade et al., 2014; Sogin et al., 2006; Vellend, 2010). Streams integrate water from multiple sources, including groundwater, storm runoff, snowmelt, and sediments, each potentially contributing distinct bacterial communities to the resulting assemblage (Besemer et al., 2013; Saup et al., 2019; Smith et al., 2018; Z. Wang et al., 2022). Consequently, variation in the relative contributions of these sources may influence community composition through stochastic recruitment of taxa, irrespective of local physicochemical conditions that drive environmental filtering or the geographic distance among sites that may impose dispersal limitation (Besemer et al., 2013; Leibold et al., 2004; Read et al., 2015; Shade et al., 2014). Terrestrial inputs may also increase particle loads, potentially enriching stream communities with particle-associated bacteria, which can exhibit higher diversity and beta dispersion than free-living bacteria (Savio et al., 2015). In addition, streams may promote local specialization because of environmental heterogeneity among sites. This may in turn contribute to the higher beta dispersion observed among stream communities and the stronger relationships between community composition and altitude, geographic distance, and physicochemical conditions (Read et al., 2015). This mechanism may be particularly relevant in our study, because stream sites spanned a broad environmental gradient ranging from alpine headwaters to lowland forests, grasslands, agricultural catchments, and urban landscapes.

In lakes, both taxonomic composition and relative abundance contributed to differences among communities across catchments. However, the greater proportion of variation explained by catchment identity using Bray–Curtis than Jaccard distances suggests that differences in the relative abundances of shared taxa contributed substantially to the differentiation of lake communities among catchments. This interpretation is supported by the dominance of a relatively small set of widespread core taxa that differ in their relative abundances. As discussed above, environmental filtering may provide one explanation for this pattern, with lake conditions favoring a common set of taxa, while physicochemical differences among lakes at the time of sampling drive variation in their relative abundances (Crump et al., 2007; Leibold et al., 2004; Lindström et al., 2005; Logue & Lindström, 2010; Vellend, 2010). These patterns align with repeatable assembly trajectories previously observed in lakes, where a dominant core microbiome emerges across catchments under stable conditions (Besemer et al., 2013; Crump et al., 2007; Niño-García et al., 2016; Papadopoulou & Lindström, 2026).

Some limitations should be considered when interpreting these findings. Catchments were sampled at different times of year, making it difficult to separate catchment differences from seasonal variation. This limitation primarily affects comparisons among catchments. Within each catchment, stream and lake samples were collected within a narrow temporal window, and differences in sampling date are unlikely to explain the consistent relationship between community structure and hydrological position. Seasonal variation may, however, help explain why Lake Pfäffikon samples clustered more closely with those from Lakes Sils and Silvaplana than with those from the nearby and hydrologically connected Lake Greifen (Figure 3A). Pfäffikon, Sils, and Silvaplana were sampled under more similar temperature conditions (Figure 1C). Given the seasonality of lake bacterial communities (Eiler et al., 2012; Stadler & Del Giorgio, 2022), clustering among lakes may therefore reflect temporal and physicochemical variation alongside spatial effects. Additional limitations concern sampling effort and detection of diversity. Sampling effort varied among catchments because of differences in stream number and length. A single sample per site prevented us from assessing within-site variability. Although rarefaction standardized sequencing depth, this may also have affected detection of rare ASVs. Replicated sampling within sites and repeated sampling across seasons could help quantify local variability and distinguish persistent community differences from temporal variation.

## Conclusion

On a landscape scale, freshwater microbial communities were structured differently across hydrological network positions in all sampled catchments. We showed that streams were overall more diverse and heterogeneous than lakes, yet had weaker between-catchment differentiation. Lakes, on the other hand, were less diverse and supported a relatively homogeneous community composition, but with distinct catchment-specific signatures. These patterns arose through a combination of shifts in the relative abundance of shared dominant taxa and turnover among less abundant taxa, with both processes more pronounced in lakes than in streams. Our findings advance our understanding of microbial biogeography in freshwater networks by showing how streams and lakes differ in the balance between community heterogeneity between sites and between-catchment differentiation.

## Acknowledgements

We thank Dr. Jean-Claude Walser for help with bioinformatics analysis. Library preparation was conducted at the Genetic Diversity Center at ETH Zurich, and sequencing was conducted at the Center of Integrative Genomics at the University of Lausanne. This work, KD and AK were supported by H2020 European Research Council, 852621.

## Artificial Intelligence disclosure

Generative artificial intelligence (AI) tools (ChatGPT, OpenAI) were used during manuscript preparation to assist with language editing and debugging code (R Wizard GPT). All analyses, statistical tests, results, interpretations, and conclusions were designed, verified, and approved by the authors. The authors take full responsibility for the accuracy and content of the manuscript.

## Data availability statement

Raw sequencing reads have been deposited in the NCBI Sequence Read Archive under BioProject accession PRJNA1450899. Analysis code and processed data will be made available through Dryad upon acceptance of the manuscript.

