## Supporting information for "Stream–lake network position structures freshwater microbial communities across multiple catchments"

**Appendix**

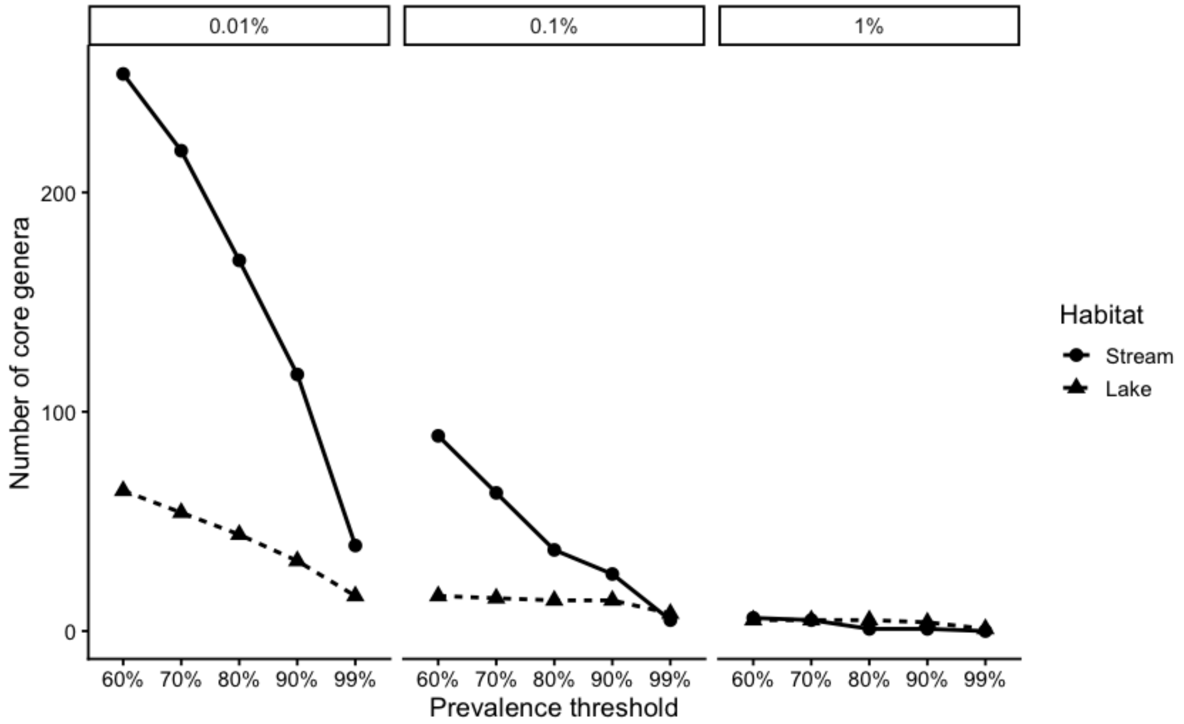


Figure S1. Sensitivity of core microbiome size to prevalence and relative abundance thresholds. The number of genera classified as core is shown across prevalence thresholds (60–99%) and relative abundance thresholds (0.01–1%) for stream and lake habitats. A prevalence threshold of 90% with a relative abundance threshold of 0.1% were chosen for delimiting core genera in this study.

Table S1: Linear mixed-effects models testing the effect of hydrological network position on alpha diversity. Observed richness was log-transformed (log1p) prior to analysis. Catchment was included as a random intercept and random slope for hydrological position.

| Response | Term | Estimate | Std_Error | DF | t_value | p_value |
|---|---|---|---|---|---|---|
| Observed (log1p) | (Intercept) | 8.655 | 0.128 | 7.1 | 67.78 | <0.001 |
| Observed (log1p) | lake_posInlet | 0.024 | 0.053 | 62.9 | 0.45 | 0.657 |
| Observed (log1p) | lake_posLake | -1.425 | 0.241 | 7.1 | -5.91 | <0.001 |
| Observed (log1p) | lake_posOutflow | -1.257 | 0.124 | 6.8 | -10.14 | <0.001 |
| Shannon | (Intercept) | 6.845 | 0.253 | 7.2 | 27.08 | <0.001 |
| Shannon | lake_posInlet | -0.152 | 0.183 | 15.3 | -0.83 | 0.418 |
| Shannon | lake_posLake | -2.598 | 0.224 | 8.7 | -11.61 | <0.001 |
| Shannon | lake_posOutflow | -2.705 | 0.377 | 24.3 | -7.17 | <0.001 |

Table S2: Tukey-adjusted pairwise contrasts of hydrological network positions for observed richness (log1p) and Shannon diversity from linear mixed-effects models including catchment-specific random intercepts and slopes. Estimates reflect differences in marginal means; SE = standard error; DF = Kenward–Roger degrees of freedom.

| Response | Contrast | Estimate | SE | DF | t_ratio | p_value |
|---|---|---|---|---|---|---|

| | | | | | | |
|---|---|---|---|---|---|---|
| Observed (log1p) | (Up-stream) - Inlet | -0.0239 | 0.0579 | 5.93 | -0.412 | 0.9743 |
| Observed (log1p) | (Up-stream) - Lake | 1.4248 | 0.241 | 6.99 | 5.906 | 0.0026 |
| Observed (log1p) | (Up-stream) - Outflow | 1.2575 | 0.124 | 6.98 | 10.106 | 0.0001 |
| Observed (log1p) | Inlet - Lake | 1.4486 | 0.24 | 6.98 | 6.033 | 0.0023 |
| Observed (log1p) | Inlet - Outflow | 1.2814 | 0.13 | 6.96 | 9.848 | 0.0001 |
| Observed (log1p) | Lake - Outflow | -0.1673 | 0.254 | 7 | -0.658 | 0.9093 |
| Shannon | (Up-stream) - Inlet | 0.152 | 0.195 | 6.12 | 0.782 | 0.8602 |
| Shannon | (Up-stream) - Lake | 2.598 | 0.233 | 6.51 | 11.132 | 0.0001 |
| Shannon | (Up-stream) - Outflow | 2.705 | 0.378 | 6.98 | 7.148 | 0.0008 |
| Shannon | Inlet - Lake | 2.445 | 0.274 | 6.52 | 8.922 | 0.0003 |
| Shannon | Inlet - Outflow | 2.552 | 0.402 | 6.96 | 6.349 | 0.0017 |
| Shannon | Lake - Outflow | 0.107 | 0.369 | 6.94 | 0.29 | 0.9907 |

Table S3: Variance components from linear mixed-effects models including stream nested within catchment. Values represent the amount of variation attributable to differences among catchments, among streams within catchments, and residual (within-group) variation. Because these values are on the scale of the response variable, they are best interpreted relative to one another rather than in absolute terms. In all cases, stream-level variance was small compared to catchment-level and residual variation.

| **Response** | **Catchment variance** | **Stream variance** | **Residual** |
|---|---|---|---|
| Temperature | 15.23 | 0.7 | 4.24 |
| pH | 0.0044 | 0.0084 | 0.064 |
| EC | 0.0206 | 0.0006 | 0.007 |
| Observed | 0.124 | 0.0055 | 0.085 |
| Shannon | 0.441 | 0.118 | 0.82 |

Table S4: PERMANOVA results (Bray–Curtis distance) testing the effect of hydrological network position on bacterial community composition, with permutations constrained within catchments.

| **Term** | **Df** | **SumOfSqs** | **$R^2$** | **F** | **p-value** |
|---|---|---|---|---|---|
| Model | 3 | 18.304 | 0.2124 | 19.596 | 0.001 |
| Residual | 218 | 67.875 | 0.7876 | | |
| Total | 221 | 86.179 | 1 | | |

Table S5: Test of homogeneity of multivariate dispersion (PERMDISP) among hydrological network positions based on Bray–Curtis distances. Significant differences in dispersion were detected among groups ($F = 78.19$, $p < 0.001$).

| **Term** | **Df** | **Sum Sq** | **Mean Sq** | **F value** | **p-value** |
|---|---|---|---|---|---|

| | | | | | |
|---|---|---|---|---|---|
| Groups | 3 | 1.3994 | 0.46647 | 78.189 | <2.2e-16 |
| Residuals | 218 | 1.3006 | 0.00597 | | |

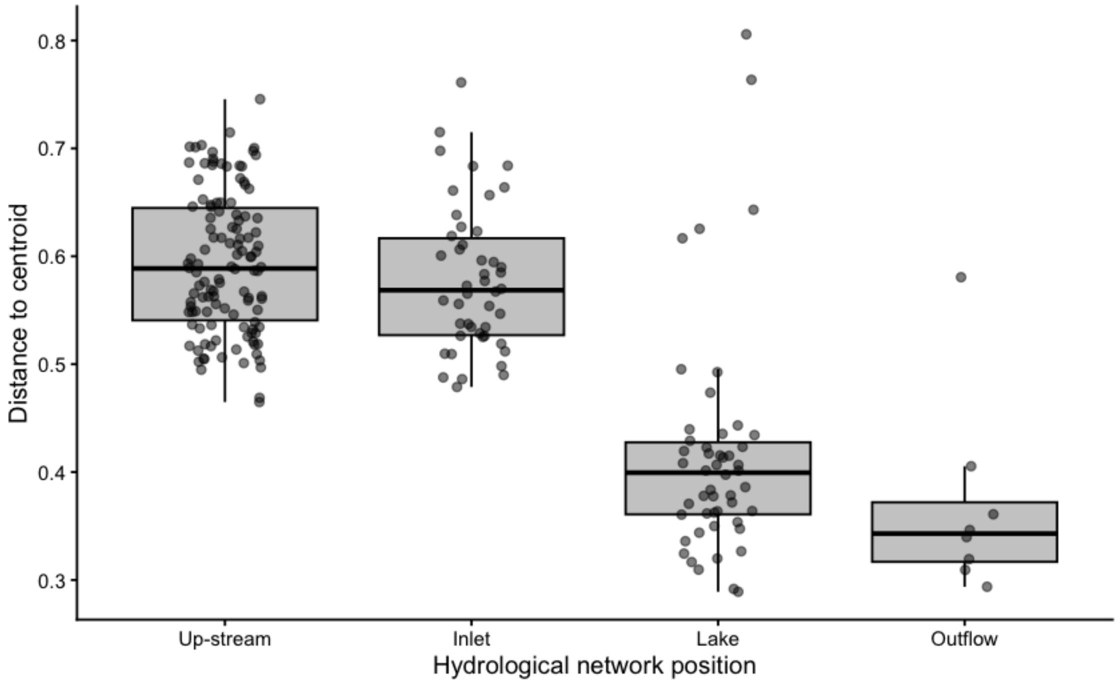


Figure S2: Distance to group centroid (PERMDISP) for hydrological network positions derived from Bray–Curtis dissimilarities. Stream habitats (Up-stream and Inlet) showed significantly greater multivariate dispersion compared to Lake and Outflow positions, reflecting higher beta-diversity heterogeneity within stream communities.

Table S6: Tukey HSD pairwise comparisons of multivariate dispersion among hydrological network positions. Differences represent contrasts in mean distance to centroid (Bray–Curtis); significant contrasts indicate greater compositional heterogeneity in stream relative to lake and outflow sites.

| **Contrast** | **Difference** | **Lower CI** | **Upper CI** | **Adjusted p-value** |
|---|---|---|---|---|
| Inlet – Up-stream | -0.0148777 | -0.0496367 | 0.01988121 | 0.6849135 |
| Lake – Up-stream | -0.1769765 | -0.2107202 | -0.1432328 | <0.001 |
| Outflow – Up-stream | -0.2238341 | -0.2968914 | -0.1507768 | <0.001 |
| Lake – Inlet | -0.1620988 | -0.2029529 | -0.1212446 | <0.001 |
| Outflow – Inlet | -0.2089564 | -0.2855579 | -0.1323548 | <0.001 |
| Outflow – Lake | -0.0468576 | -0.1230038 | 0.02928865 | 0.3846538 |

Table S7: Mantel correlations between community dissimilarity (Bray–Curtis) or environmental dissimilarity (Euclidean distance of rank-transformed variables) and spatial gradients (geographic distance and altitude) in stream and lake habitats. Mantel tests were calculated using only between-catchment sample pairs**.**

| **Habitat** | **Predictor** | **Response variable** | **Mantel r** | **p-value** |
|---|---|---|---|---|
| Lake | Altitude | Community | 0.108 | 0.035 |
| Lake | Altitude | EC | 0.473 | 0.001 |
| Lake | Altitude | Temperature | 0.292 | 0.001 |
| Lake | Altitude | pH | 0.082 | 0.035 |
| Lake | Distance | Community | 0.192 | 0.011 |
| Lake | Distance | EC | 0.490 | 0.001 |
| Lake | Distance | Temperature | 0.312 | 0.001 |
| Lake | Distance | pH | 0.028 | 0.559 |
| Stream | Altitude | Community | 0.437 | 0.001 |
| Stream | Altitude | EC | 0.591 | 0.001 |
| Stream | Altitude | Temperature | 0.591 | 0.001 |
| Stream | Altitude | pH | 0.051 | 0.016 |
| Stream | Distance | Community | 0.557 | 0.001 |
| Stream | Distance | EC | 0.474 | 0.001 |
| Stream | Distance | Temperature | 0.339 | 0.001 |
| Stream | Distance | pH | 0.010 | 0.731 |

Table S8: Mantel test results showing relationships among environmental dissimilarity (Euclidean distance of rank-transformed temperature, electrical conductivity, and pH), community dissimilarity (Bray–Curtis), geographic distance, altitude, and sampling-date differences across stream and lake habitats. Mantel tests were calculated using only between-catchment sample pairs.

| **Habitat** | **Predictor** | **Variable** | **Mantel r** | **p-value** |
|---|---|---|---|---|
| Stream | Distance | Environmental dissimilarity | 0.396 | 0.001 |
| Stream | Distance | Community dissimilarity | 0.557 | 0.001 |
| Stream | Altitude | Environmental dissimilarity | 0.595 | 0.001 |
| Stream | Altitude | Community dissimilarity | 0.437 | 0.001 |
| Stream | Sampling date | Environmental dissimilarity | 0.151 | 0.001 |
| Stream | Sampling date | Community dissimilarity | 0.001 | 0.971 |
| Stream | Community | Environmental dissimilarity | 0.478 | 0.001 |
| Lake | Distance | Environmental dissimilarity | 0.411 | 0.001 |
| Lake | Distance | Community dissimilarity | 0.192 | 0.011 |
| Lake | Altitude | Environmental dissimilarity | 0.401 | 0.001 |
| Lake | Altitude | Community dissimilarity | 0.108 | 0.035 |
| Lake | Sampling date | Environmental dissimilarity | 0.114 | 0.008 |
| Lake | Sampling date | Community dissimilarity | 0.246 | 0.002 |
| Lake | Community | Environmental dissimilarity | 0.369 | 0.001 |

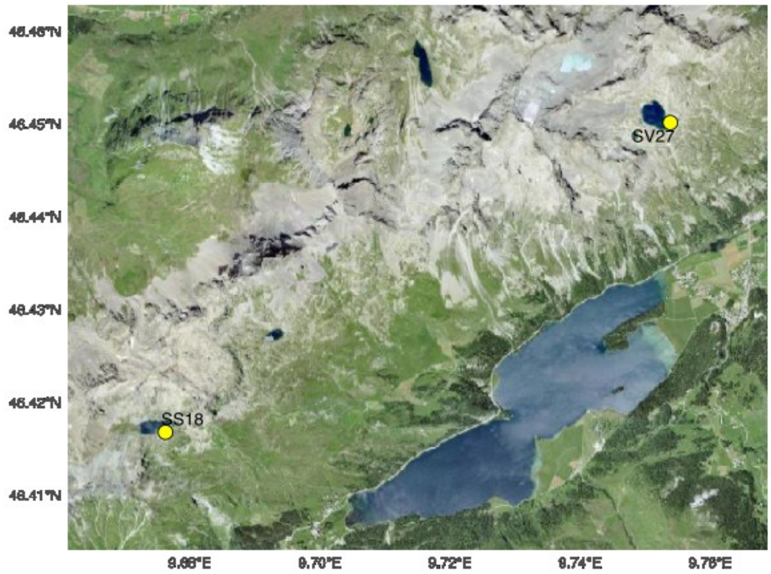


Figure S3: Geographic location of SS18 and SV27 sampling sites shown on a satellite basemap, showing that the sampled streams originate from small glacial lakes.

Table S9: PERMANOVA results testing the effect of catchment on bacterial community composition (Bray–Curtis dissimilarity) within stream and lake habitats, partitioned into full, core, and peripheral genera.

| **Habitat** | **Dataset** | **Numer of genera** | **$R^2$** | **F statistic** | **p-value** |
|---|---|---|---|---|---|
| Stream | Full | 1078 | 0.23 | 6.71 | 0.001 |
| Stream | Core | 26 | 0.17 | 4.61 | 0.001 |
| Stream | Peripheral | 1052 | 0.26 | 7.64 | 0.001 |
| Lake | Full | 917 | 0.58 | 9.91 | 0.001 |
| Lake | Core | 14 | 0.57 | 9.62 | 0.001 |
| Lake | Peripheral | 903 | 0.64 | 12.79 | 0.001 |

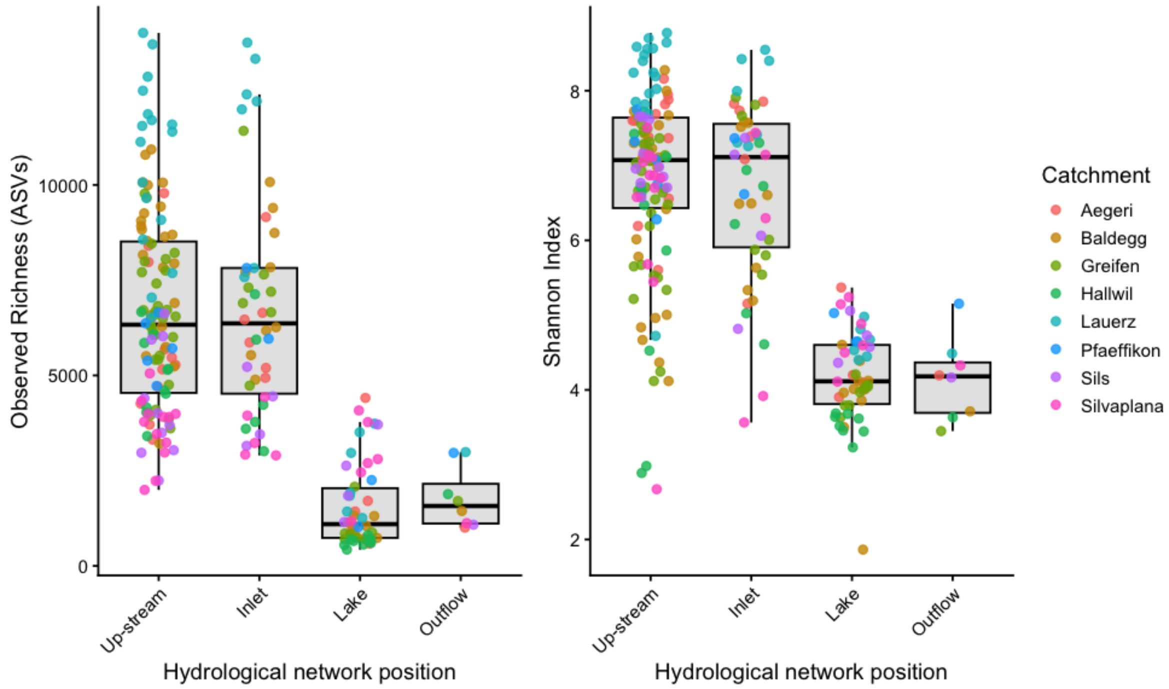


Figure S4: Alpha diversity across hydrological network positions. Boxplots show the distribution of observed ASV richness (left) and Shannon diversity (right) across samples, with individual points colored by catchment.

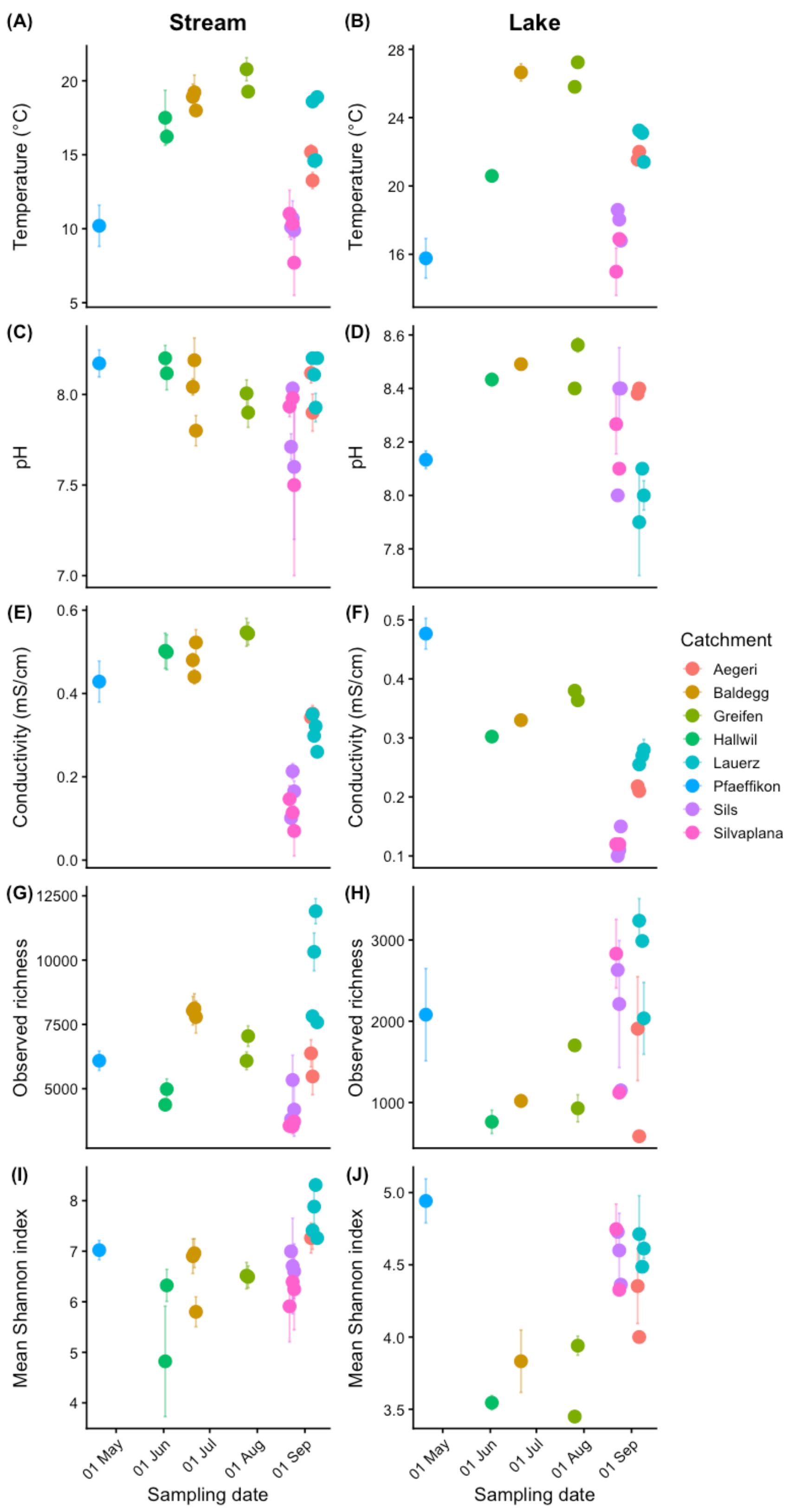


Figure S5: Temporal variation in environmental conditions and bacterial alpha diversity. Mean temperature (A–B), pH (C–D), electrical conductivity (E–F), observed richness (G–H), and Shannon diversity (I–J) are shown for stream (left column) and lake (right column) communities. Points represent catchment-level means of groups of samples collected on different days, and error bars indicate ±1 standard error. Colors denote catchments.

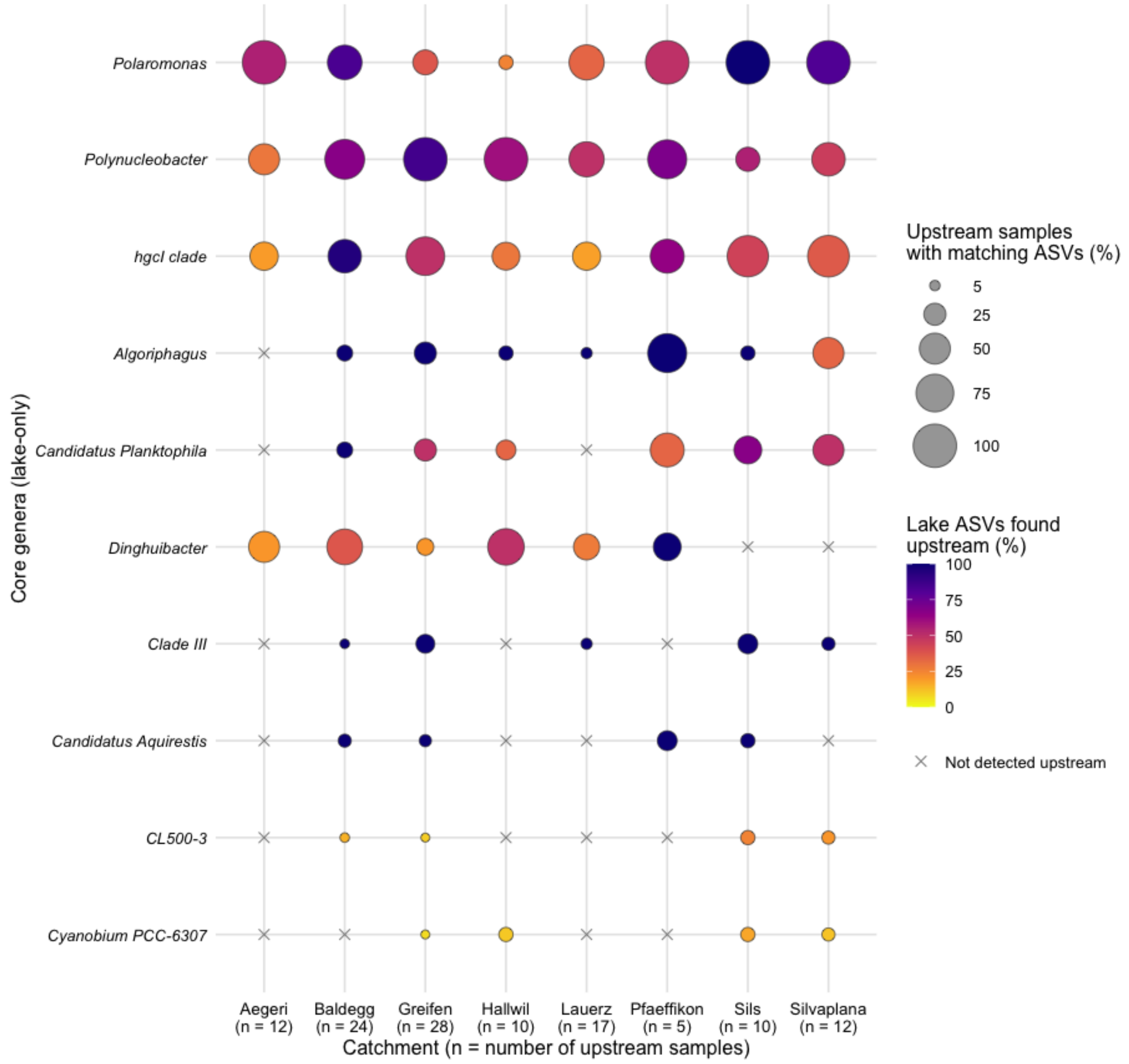


Figure S6: Bubble matrix showing that lake-core bacteria are also detected upstream. Rows show all genera that met the core criteria in the lake but not in stream communities. The size of each bubble represents the percentage of ASVs from one lake-core-only genus (rows) that were detected in both the lake and at least one upstream sample within the same catchment (columns). Bubble size represents the percentage of upstream samples in a given catchment containing at least one ASV from a given lake-core-only genus. Crosses indicate no matching ASVs. Detection required at least five reads per ASV in an individual sample. Inlets were excluded from the comparison.
